# Combinatorial analysis of *Saccharomyces cerevisiae* regulatory elements

**DOI:** 10.1101/777763

**Authors:** N. Dhillon, R. Shelansky, B. Townshend, M. Jain, H. Boeger, D. Endy, R.T. Kamakaka

**Affiliations:** Department of MCD Biology, 1156 High Street, University of California, Santa Cruz, CA 95064 USA; Department of Bioengineering, Stanford University, Stanford, CA 94305, USA; Department of Biomolecular Engineering, 1156 High Street, University of California, Santa Cruz, CA 95064 USA

## Abstract

Gene expression in *Saccharomyces cerevisiae* is regulated at multiple levels. Genomic and epigenomic mapping of transcription factors and chromatin components has led to the definition and delineation of various regulatory elements. Enhancers, promoters, 5’ untranslated regions (5’UTR) and transcription terminators/3’ untranslated regions (3’UTR) have all been defined. However, the specific contributions of each of these features as part of a regulatory unit and the functional communications between these regulatory elements remains under explored.

We built a combinatorial library of 26 different enhancers, core promoters, 5’UTRs and transcription terminators/3’UTRs. This library was analyzed with respect to gene expression in order to better understand the interactions between different regulatory elements. In the process we developed new methods to estimate the contribution of individual regulatory parts from just a few simple measurements. Our data show that different pairs of regulatory parts follow specific interaction rules affecting overall activity either positively or negatively. We find that while enhancers are the initiators of gene activity, core promoters modulate the levels of enhancer mediated expression. Cluster analysis based on expression show that TATA-box containing core promoters appear to increase enhancer-driven transcription to a greater extent than TATA-less promoters. Principal component analysis highlight outliers and suggest differences in mechanisms of regulation. These results provide a system to characterize regulatory elements and use these elements in the design of synthetic regulatory circuits.

## Introduction

Gene expression in eukaryotes is regulated at multiple levels including at the level of transcription, post-transcriptional mRNA control and translational control. Regulatory elements have been identified and categorized based on their effects on gene expression at the different levels of regulation. Mutational analysis as well as genomic and epigenomic mapping of proteins has led to the definition and delineation of enhancers, promoters, 5’ and 3’ untranslated regions as well as transcriptional terminators. The regulatory landscape is compartmentalized into enhancer elements that allow increased expression of mRNA in response to signals and promoter sequences that function as binding sites for general transcription factors and sites of transcription initiation. The 5’ untranslated region is involved in the association of the mRNA with the ribosome while the 3’ untranslated region is involved in mRNA stability and turnover. Regulation of these elements is mediated by sequence specific DNA binding proteins that recognize their cognate binding sites in DNA as well as RNA binding proteins that mediate their effects via interactions with mRNA.

The identification of DNA binding proteins and the mapping of protein binding motifs across the genome has been helpful in understanding the regulation of gene expression (Badis et al., 2009; Gerstein et al., 2012; Johnson et al., 2007; Jolma et al., 2013; Ren et al., 2000; Rhee and Pugh, 2012; Venters et al., 2011; Zhu et al., 2009) but determining the relative contribution of each regulatory element in gene regulation has proven to be labor intensive and time consuming. Initial studies on gene regulation investigated regulatory elements within single genes via directed mutagenesis of specific regulatory elements such as enhancers and core promoters coupled with measurements of gene activity (Hahn and Young, 2011; Maniatis et al., 1987; Smale and Kadonaga, 2003). These analyses gave way to saturation mutagenic studies of single regulatory elements against a background of unmodified elements (Lubliner et al., 2015) as well as approaches where entire regulatory domains of individual genes were subjected to saturation mutagenesis coupled to experiments studying the effects of these mutations on mRNA and protein synthesis (Patwardhan et al., 2009). While very valuable, these approaches were resource consuming and were restricted to specific genes.

Alternative approaches for investigating the complex regulatory landscape involved generating random combinations of individual sequence elements to achieve a desired expression level and then deconvoluting the identity of the regulatory elements to decipher their individual role in the process (Grossman et al., 2017; Kosuri et al., 2013; Mutalik et al., 2013a; Mutalik et al., 2013b). Such combinatorial analyses of regulatory elements have begun to allow the investigation of the effects of combining different regulatory elements.

We set out to characterize a set of regulatory elements in the yeast *Saccharomyces cerevisiae*. We delineated a gene into four regulatory fragments: enhancer, core promoter, 5’UTR and 3’UTR/terminator and built a library of each of these fragments from 26 different genes. We then used a directed approach to rejoin the fragments in the correct order thus creating complete synthetic genes with parts chosen at random from these 26 different genes. This combinatorial library was fractionated on the basis of expression levels of a fluorescent reporter gene using flow cytometry and cell sorting. The reporter gene with its associated regulatory elements from the fractionated cells were sequenced using next generation nanopore sequencing to identify the regulatory elements. These data were analyzed to identify the role of each regulatory fragment in mediating specific levels of gene expression.

We validated this combinatorial approach by building a matrix where nine different enhancer sequences were combined with nine different promoter elements and the resulting 81 constructs were analyzed for expression under varying environmental and mutagenic conditions.

## Results

### Fragmentation of the Regulatory Space and Combinatorial Library Preparation

The standardization of biological parts and their characterization under varying genetic background, media, growth and environmental conditions is necessary in order for these parts to be routinely mixed and matched (Endy, 2005). Transcription of protein coding genes in yeast is primarily mediated by regulatory sequences located upstream of the start of transcription. These sequences have an upstream activating enhancer sequence (UAS) that binds sequence specific transcription factors which in turn direct transcription from the core promoter made up of the TATA box and initiator element that are bound by the general transcription factors. Often the promoter and UAS enhancer are conflated together but, in this manuscript, we will use the term core promoter to refer to the DNA elements that are bound by the general transcription factors and the polymerase and the term enhancer to refer to the UAS (Blazeck and Alper, 2013).

To initiate this study, we chose 26 different yeast genes based on their expression profiles in glucose (Nagalakshmi et al., 2008; Rhee and Pugh, 2012; Xu et al., 2009) (Figure 8). We demarcated their regulatory space using different genomic databases (Supplementary figure 1). We delineated the 3’UTR and transcription terminator of these genes based on the functional characterization of transcription terminators (Yamanishi et al., 2013) and identified the 5’UTR based on RNA-seq analysis of transcription in glucose (Nagalakshmi et al., 2008; Rhee and Pugh, 2012; Xu et al., 2009). Annotation of the core promoter of these yeast genes was based on ChIP-Seq mapping of TBP and other general transcription factors (Rhee and Pugh, 2012). Half of these genes have a TATA box while the rest are considered “TATA-less” or were unannotated (Rhee and Pugh, 2012). The upstream enhancer sequences of the genes were initially identified in silico using the ChIP-Seq data of various sequence specific transcription factors (Venters et al., 2011) but we decided to take the entire DNA sequence from the core promoter to the start or stop codon of the upstream gene to be the UAS enhancer fragment. To confirm the validity of the delineation, we analyzed the chromatin architecture of the regulatory region of the 26 different genes. Using ATAC-seq data (Schep et al., 2015) we mapped highly accessible regions, which have previously been shown to occur at protein binding sites. We also mapped TBP binding, histone occupancy and nucleosome positions at these genes to identify promoters and enhancers (Supplementary figure 1) (Brogaard et al., 2012; Dion et al., 2007; Hamdani et al., 2019; Rhee et al., 2014; Rhee and Pugh, 2012). Consistent with our delineation of the regulatory fragments, we find that TBP binding coincides within the delineated core promoters, while accessible sites based on ATAC-seq map primarily to the enhancer fragments and nucleosome depleted regions map to both enhancers and core promoters as well as transcription terminators.

Each of the delineated regulatory fragments from the 26 different genes were PCR amplified with primers such that they contained two different type IIS restriction enzyme sequences. The 104 (26×4) PCR fragments were amplified from yeast genomic DNA and cloned into pYTK001 using BsmBI and the Golden Gate cloning methodology (Lee et al., 2015). We were able to clone 103 out of the 104 expected plasmids but were unable to clone the *HIS3* promoter fragment. In addition to the individual elements we have also constructed other combinations of elements (which were then used to generate constructs used in figures 8-11). The 103 parts plasmids were then used to create a combinatorial library such that the fragments would combine in a directed but random manner into a *ARS/CEN/URA3* plasmid, a derivative of pYTK096 (Lee et al., 2015). The large combinatorial ligation reaction was used to transform *E. coli* STBL4 cells by electroporation and the transformants were selected for Kanamycin resistant colonies. A 1x coverage of all possible combinations would result in 439,400 clones and we obtained approximately 600,000 *E. coli* colonies. Based on random sampling, we would expect 74% of the possible clones to be present in this *E. coli* transformed library at least once (Clarke and Carbon, 1976). The colonies were scrapped from the plates and pooled. The plasmid DNAs were then isolated from these cells and purified on a cesium chloride gradient.

### Measurement of gene activity

The purified library was transformed into W-303 yeast cells (ROY5634) by electroporation and transformants were selected on YMD-uracil plates. This strain contains a fluorescent protein mTagEBFP2-2 under the control of the *RPL18b* promoter (The published yeast mTagEBFP2 sequence (Lee et al., 2013) does not generate a fluorescent protein (data not shown). We therefore integrated a codon optimized version of the mammalian mTagEBFP2 protein (Subach et al., 2011) that we refer to as mTagEBFP2-2). Approximately 200,000 yeast transformants were scrapped from the YMD-uracil plates and frozen (as the unamplified yeast library).

Cells containing the library were grown under selection to log phase, pelleted and resuspended into 250ul 1xPBS and 1%BSA at a concentration of 1 OD/ml, filtered through a Nitex mesh and cells were sorted using a fluorescence assisted cell sorter (FACS) into four bins based on expression of both mRuby2/EBFP2-2. The gates for fluorescent cell sorting were based on various control strains. Prior to sorting the library, we analyzed four different transformants, a strain that did not express mRuby2 or mTagEBFP2-2, a strain that only expressed mTagEBFP2-2, a strain that only expressed mRuby2 and a strain that highly expressed mRuby2 along with mTagEBFP2-2. We sorted the yeast cells containing the library based on the expression level of mRuby2 compared to the constitutive expression of mTagEBFP2-2. We gated the fractions into 4 expression categories (no expression, low, medium and high expression) and collected 24,540,211 cells (Figure 1). 63% of the cells were in the no expression fraction, 26% were in the low expression fraction, 8% were in the medium expressing fraction and 3% were in the high expressing fraction

**Figure 1:**
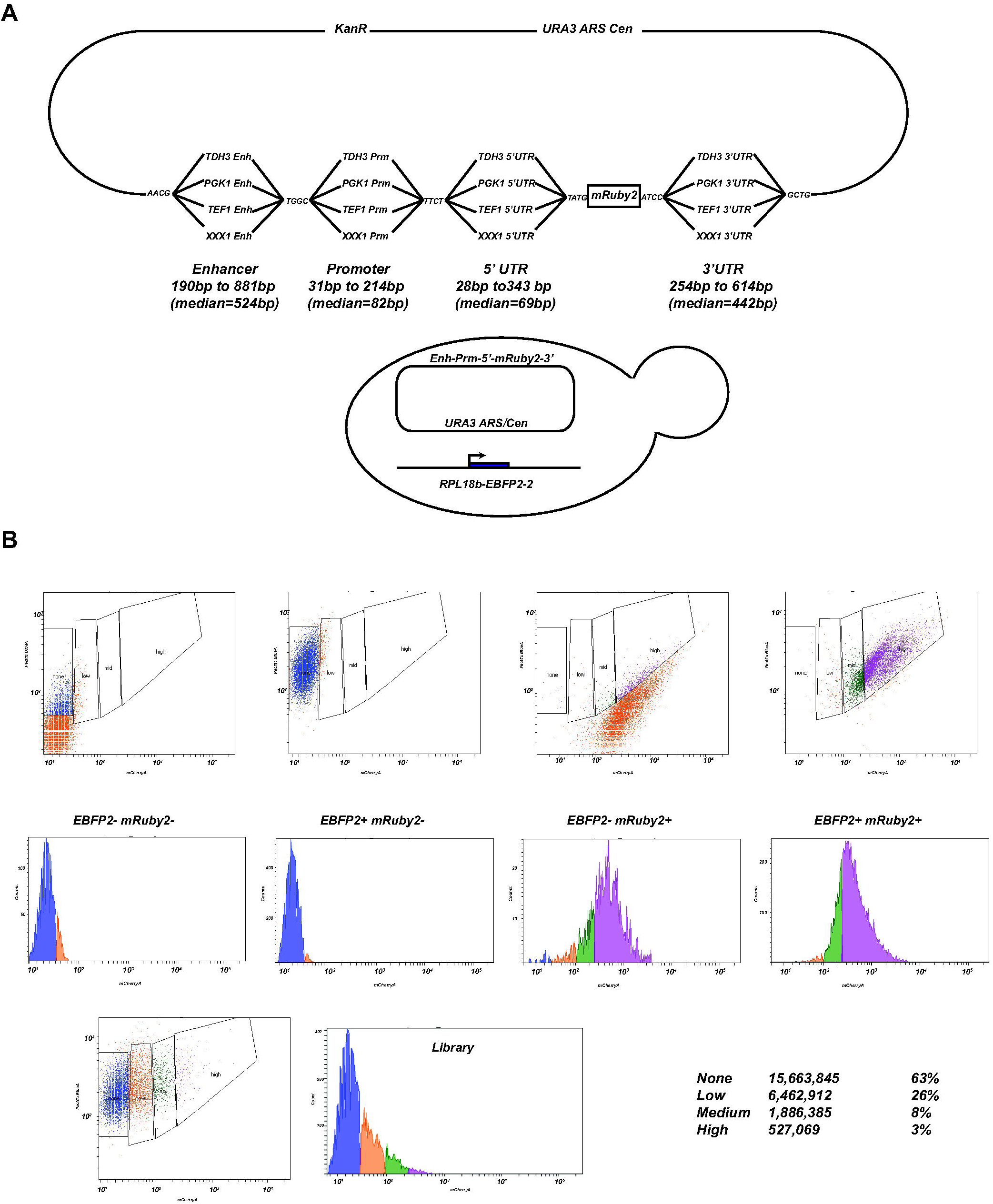
Library construction and sorting A Schematic of the assembly of the regulatory elements to recreate a functional gene cassette with mRuby2 as a reporter gene. The constructed library was used to transform yeast cells expressing EBFP2-2 under the control of the *RPL18b* promoter. B Cytometry traces for various control strains used to generate gates for cell sorting as well as the sorted bins of the library based on Red:Blue fluorescence ratio. Each sorted bin had different number of cells.

After growth of the cells for a further 12h at room temperature, genomic and plasmid DNA were isolated using a Yeastar Genomic DNA Kit (Zymo Research). The entire insert (including mRuby2) was PCR amplified with primers flanking the expression cassette. Barcodes were ligated to the amplified fragments to distinguish the four sorted pools and the PCR products were subsequently sequenced using an Oxford Nanopore MinION sequencer.

We obtained 1,662,773 reads from the 4 sorted fractions that mapped to the 26 gene fragments. Seventy three percent of the mapped reads had all four DNA regulatory fragments in the correct order while the remainder lacked one or more fragment either due to inappropriate joining during ligation or to our inability to unambiguously assign an identity to the fragment after sequencing. Of the total mapped reads, 61% were from the no mRuby2 expression fraction, 19% were from the low expression fraction, 5% were from the medium mRuby2 expression fraction and 14% were from the high mRuby2 expressing fraction.

We next analyzed the distribution of reads across the different regulatory fragments for the data from the sorted cells. The distribution of these regulatory fragments is shown (Figure 2) and indicate that most of the expected regulatory fragments are present in the clones sequenced from yeast cells. Several fragments were either absent or had only a few reads associated with them-*TEF1, PHO5, GLK1, HIS3* and *ADH1* enhancer fragments, *ICL1* and *ACT1* 5’UTR, and the *PGI1, HIS3, ACO1* and *TEF1* 3’UTR fragments. As expected, we also did not find any reads for the *HIS3* promoter.

**Figure 2:**
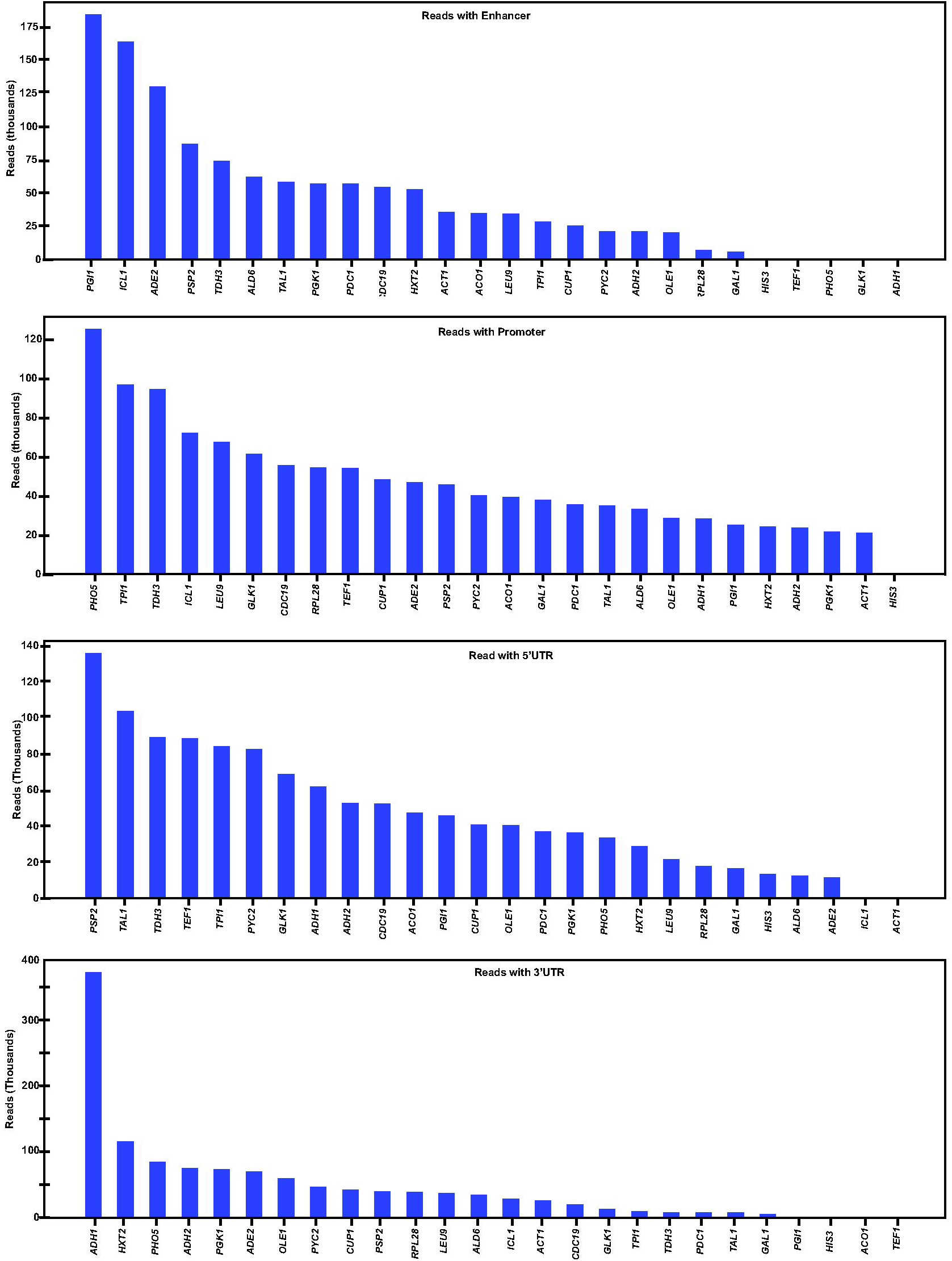
Sequence Analysis of sorted bins Number of reads for each individual regulatory element from the sorted library.

### Regulatory element correlation with transcription activity

The number of total reads in each of the four expression fractions parallels the number of cells sorted into each expression fraction. The ratio of a specific fragment in each fraction was determined by normalizing the number of reads for a fragment in each sorted fraction by the sum of the reads for that fragment across all four fractions. We then pooled the reads by regulatory elements and ordered each fragment by the ratio present in each fraction as percent of the total (Figure 3). Enhancer fragments for the inducible genes *GAL1, ADH2, CUP1 and ICL1* are present almost exclusively in the inactive non-expressing fraction while the *TDH3* and *RPL28* enhancers are enriched in the high and medium expressing fractions. This is consistent with expression analysis of yeast cells growing in glucose (Nagalakshmi et al., 2008; Rhee and Pugh, 2012; Xu et al., 2009).

**Figure 3:**
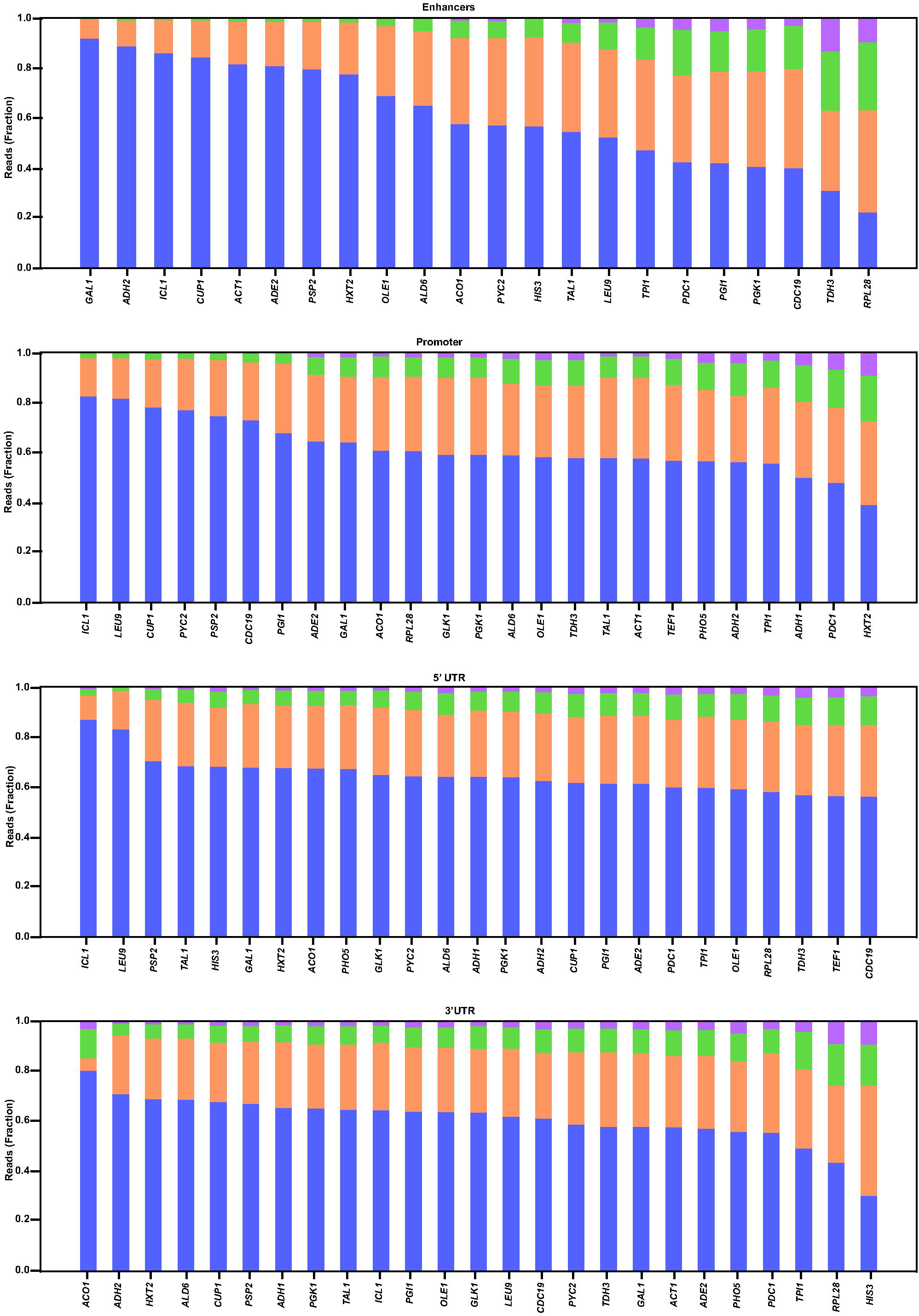
Stacked histograms of the percent of each regulatory element (Enhancer, promoter, 5’UTR and 3’UTR/terminator) present in the four sorted bins. Blue bars are from non-expressing bin, orange bars are from low expressing bin, green is from medium expressing bin and purple is from the high expressing bin.

However, when the same analysis was done with the promoter fragments, we find interesting discordance. *PGI1* and *CDC19* are both highly expressed in glucose containing media, but their promoter fragments are present to a greater extent in the non-expressing fraction. In addition, promoters of genes not active in glucose containing media such as *HXT2, PHO5* and *ADH2* are present to a greater extent in the highly expressing fraction.

The 26 different 5’UTR fragments have similar distributions across the four sorted fractions suggesting that they play a smaller role in regulating levels of gene expression.

### Functional Interactions between Regulatory elements

We next calculated the mean expression controlled by a regulatory fragment by taking a weighted average of the mean fluorescence for each sorted cell fraction, as determined by FACS, where the weights were determined by the number of reads for each fragment within each of the four expression fractions. Thus, the minimum expression that could be achieved by a regulatory fragment would occur if that fragment was solely present in the no-expression FACS fraction. Similarly, maximum expression by a regulatory fragment would be achieved if a regulatory fragment was solely present in the highly-expressing FACS fraction. Since, many regulatory fragment combinations had low sequencing depth, to increase statistical power, we pooled reads from individual regulatory elements. We then calculated the distribution of expression mediated by one individual regulatory element (e.g. enhancers) with respect to a second regulatory element (e.g. promoters). The expression mediated by each enhancer fragment was therefore determined when paired with the 26 different promoters. In figure 4 (top panel) we show the expression data for each individual enhancer with respect to the 25 different promoters (each promoter is represented by a single dot; the black dot is that enhancer with its native promoter). The enhancers were ordered based on mean expression levels. For the highly expressing enhancers, certain promoters enabled higher expression while other promoters reduced expression such that the levels of expression varied significantly between different promoter fragments. This suggests that core promoters modulate levels of enhancer mediated transcription. The level of expression mediated by a given enhancer were also influenced by the different 5’UTRs and the 3’UTRs though the variance in expression was not as large as those seen for the promoter fragments. The same effect is observed when one focuses on promoter elements (data not shown). The greatest variation in the expression from a given promoter was observed when that promoter was paired with the different enhancers while the UTRs alter promoter function to a lesser degree.

**Figure 4:**
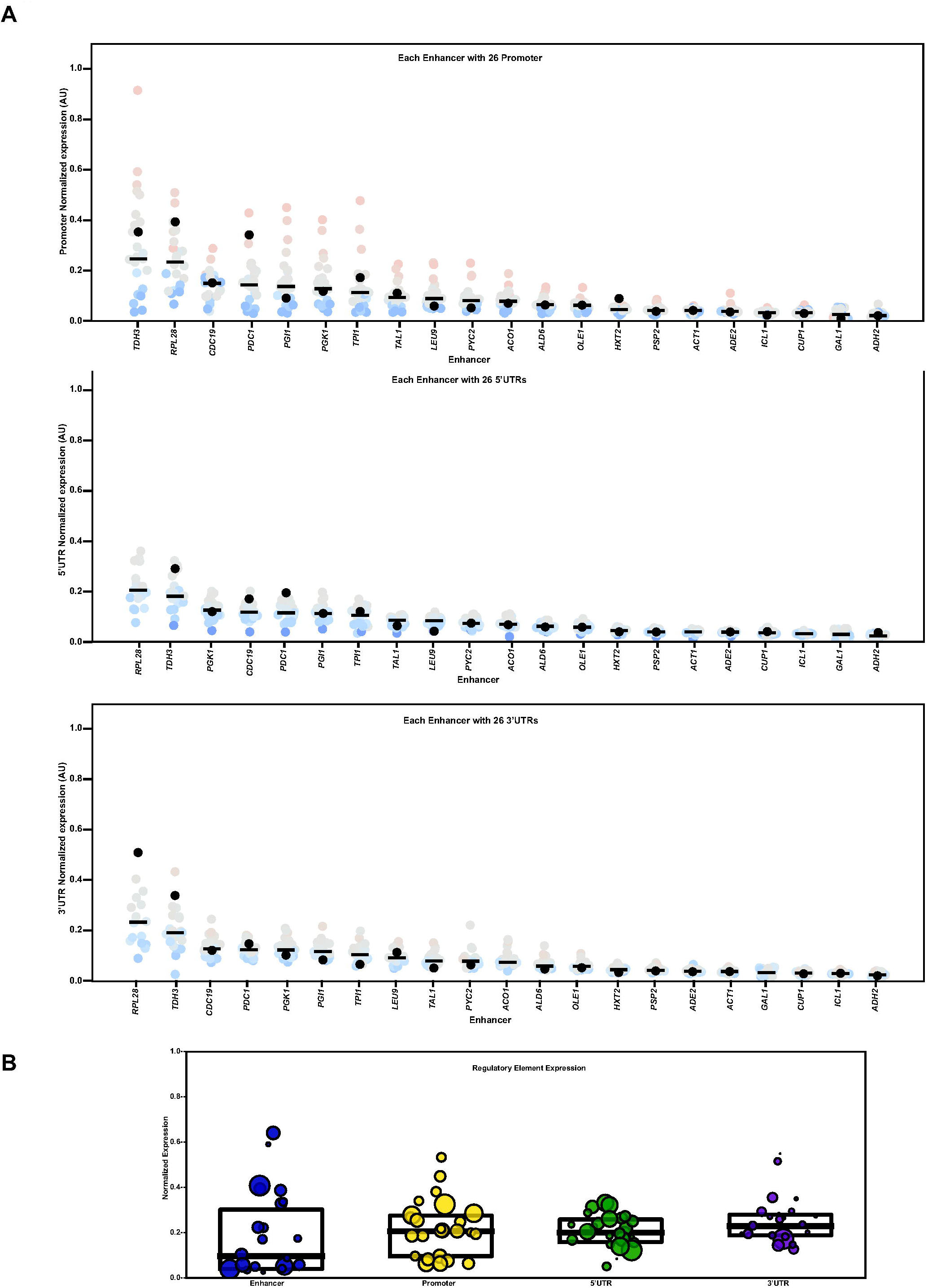
Expression analysis of pairwise combinations of regulatory elements. A The mean expression of a regulatory fragment was generated by taking a weighted average of the fluorescence for each expression category (determined by FACS) times the number of reads of each fragment within each of the four sorted bins. Pairwise comparison of Enhancers with Promoters, Enhancers with 5’UTRs and Enhancers with 3’UTRs are shown. Each enhancer is shown on the X-axis. Each dot in the box plot represents one specific regulatory element (promoter, 5’UTR or 3’UTR). B A box plot showing the normalized expression of all 26 regulatory elements are shown

We analyzed the correlation between enhancer activity and promoter activity. There was no correlation between the rank order of enhancer activity and the rank order of promoter activity. This is likely due to the fact that the inducible genes we analyzed here are expected to be inactive in YMD-uracil media. When their native promoters are separated from their cognate enhancers and the promoters are ectopically paired with other enhancers, then these promoter’s innate ability to foster high expression manifests itself. It is for this reason that *HXT2* and *ADH2* enhancers are inactive but their promoters are some of the strongest promoters.

To better quantify the ability of promoters to alter the expression mediated by enhancers we reasoned that how far one regulatory fragment such as a specific enhancer shifts another regulatory fragment such as a promoter from its mean is an indication of the strength of that element. We calculated the z-score as a metric of strength for each regulatory fragment based on the number of standard deviations from the mean that they drag a coupled fragment (Figure 5). We reasoned that this would generate a value representing how over or under-expressed one regulatory element was when paired with a second regulatory component. The data suggest that strong enhancers move promoter elements 2-3 standard deviations from their mean while weaker enhancers move promoters to a lesser degree. Similarly, strong promoters move expression of an enhancer element by 1-2 standard deviations. However, 5’UTRs move a enhancer or a promoter from the mean to a lesser degree with a uniform z-score distribution around 1. Similarly, there is less variance when comparing the effects of 5’UTRs on 3’UTRs and vice versa.

**Figure 5:**
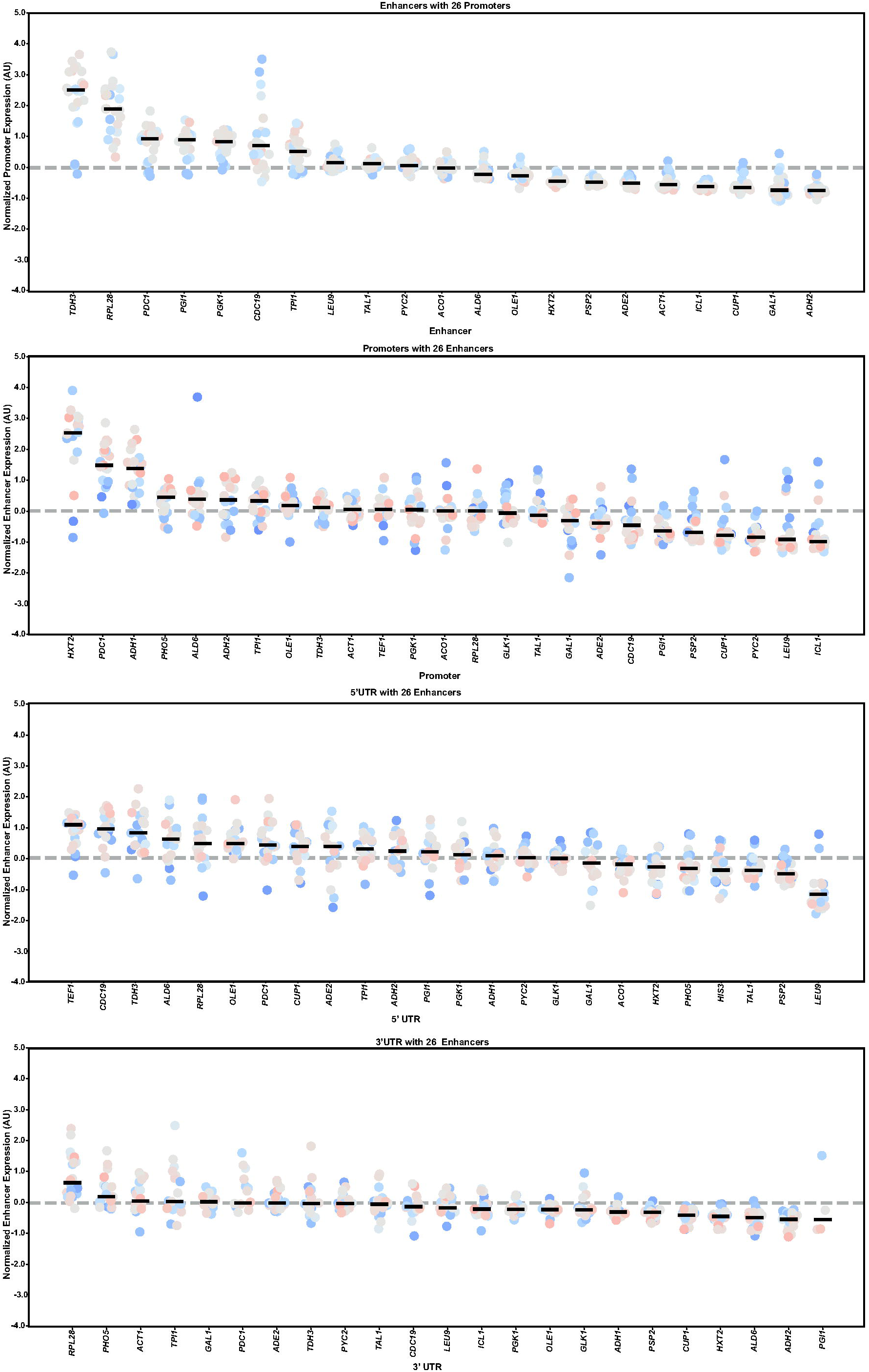
Z-scores of pairwise combinations of regulatory elements as a metric of strength. Calculated Z-scores of each pairwise combination of regulatory elements was generated and plotted. Individual regulatory elements are shown on the X-axis. Each dot in the box plot represents another specific regulatory element (enhancer, promoter, 5’UTR or 3’UTR).

### Heat maps and clustering

The same pairwise comparisons can also be depicted as heat maps and clustered using hierarchical clustering based on expression levels to identify elements with similar expression profiles. Clustering of enhancer-promoter pairs shows distinct expression profiles. These clustering profiles cannot be explained simply by the presence of specific transcription factor binding sites in the enhancer such as Reb1, Gcr1, Rap1 or Abf1. Nor are they explained by the presence of highly accessible regions as denoted by ATAC-seq or the presence and size of NFRs as determined by histone ChIP-Seq and nucleosome mapping. To some extent, promoters characterized as “TATA-less” cluster as weak promoters while TATA containing promoters cluster together as strong promoters (Figure 6).

**Figure 6:**
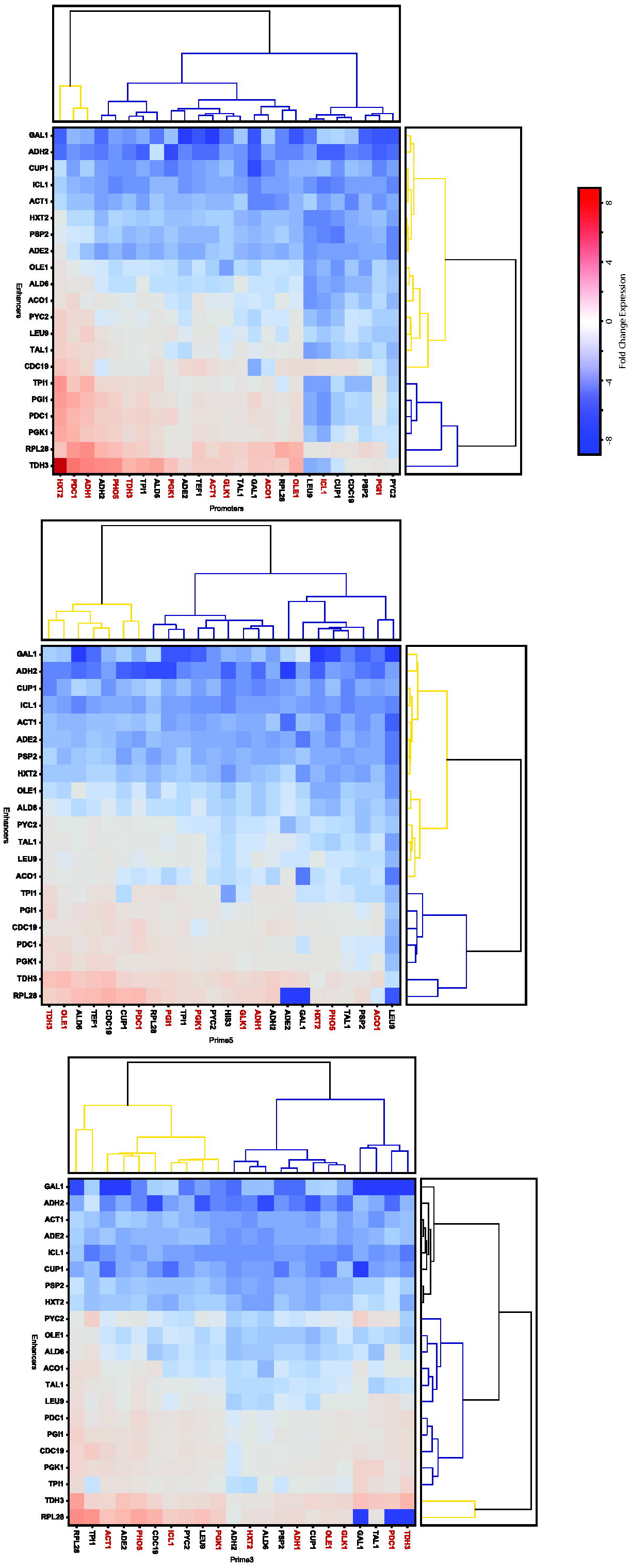
Heat maps and clustering to identify relationships between pairwise regulatory elements. The calculated expression values of each pairwise combination of regulatory element was plotted as a heat map and the values were used to generate clusters based on expression levels. TATA box containing core promoters are labeled in red.

**Figure 7:**
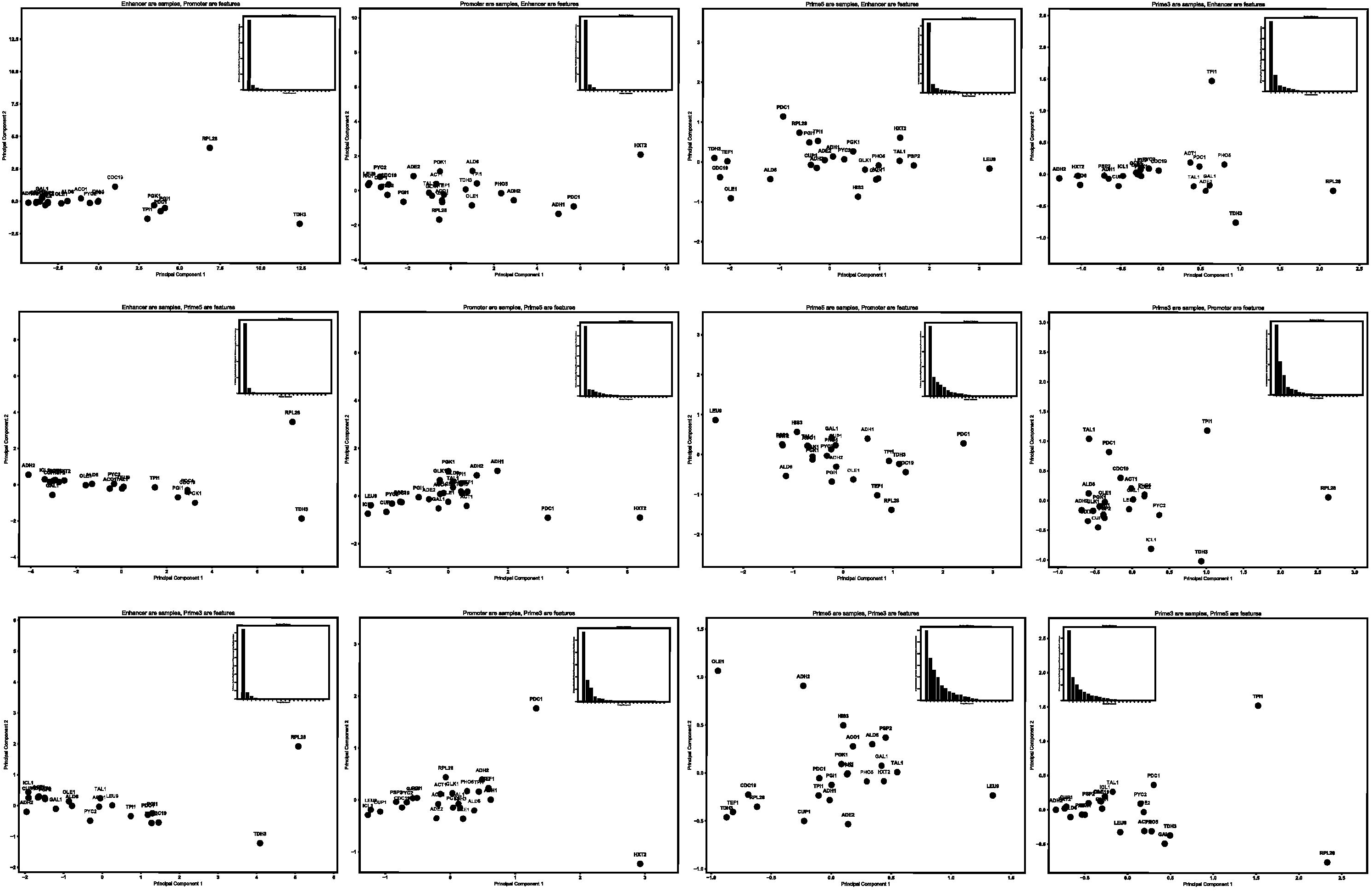
Principal component analysis of pairwise combinations of regulatory elements. The score plot displays each sample with respect to the first two principal components and was used to determine the relationship among the samples. The proportion of variance present within each principal component is plotted as an inset.

**Figure 8:**
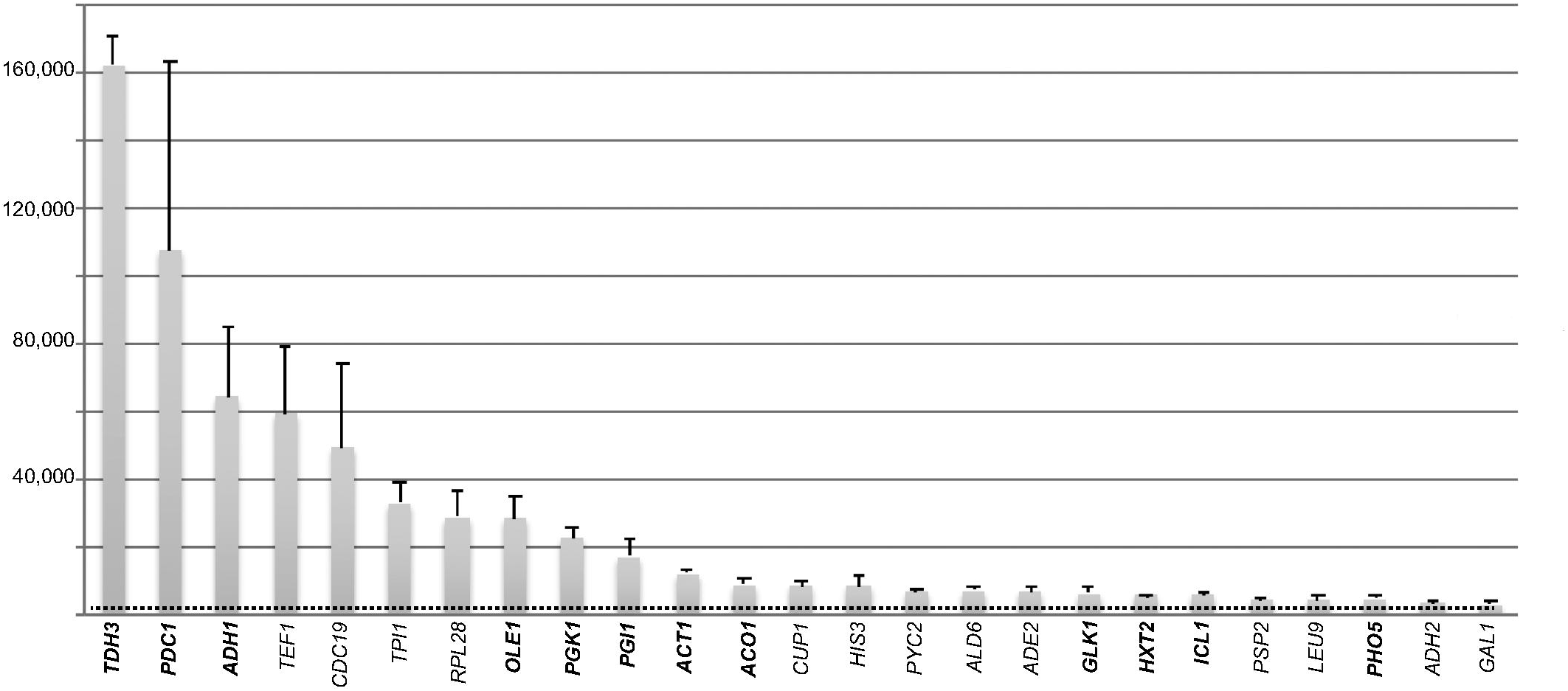
Expression of each of the native regulatory elements. The native Enhancer-promoter-5’UTR of each of the genes were fused to a Venus reporter gene and a PGK1 3’UTR and expression of the Venus reporter was measured using a fluorometer and plotted.

### Principle component analysis highlights regulatory element outliers

To depict the relationships between regulatory fragments we used Principal Component Analysis (PCA). PCA reduces dimensionality but retains the variance, interactions and correlations between samples within large datasets thus highlighting the sources of variation and the distribution of the data. We therefore performed PCA using one regulatory element as samples and another regulatory element as features.

A PCA using enhancers as samples and promoters as features, effectively distributed enhancers in promoter space. This analysis found that one predominant axis of variation across the 26 different promoters explained ∼90% of the total variance. Plotting the 26 enhancers across the first two principal components shows no distinct clusters, rather a gradient emerges. This distribution mirrors the rank in median z-scores. We therefore infer that PCA1 reflects the ability of an enhancer to amplify expression-enhancer strength. Since similarities in the data are correlated with distance in the projection space (defined by the PCA) the analysis also allows us to identify outliers and the *RPL28* enhancer occupies a distinct position on PCA2, while *TDH3* occupies a distinct position along PCA1. This suggests that these two enhancers communicate with various promoters in a manner that is distinct and different from the other elements. PCA done pairwise across the other elements, identified outliers unique to specific pairwise interactions.

The same analysis with promoters as samples and enhancers as features, showed that one principal component explains ∼90% of the total variance and PCA1 likely reflects promoter strength. Interestingly, PCA1 for 5’UTRs and 3’UTR’s explains only ∼60% of the total variance suggesting a more complex regulatory relationship between the UTRs and the other elements. The *LEU9* 5’UTR appears to be an outlier in its ability to repress expression while the *RPL28* and *TPI1* 3’UTR is an outlier that is able to enhance expression.

### 9×9 Matrix of Enhancers and Promoters

To validate results seen by FACS and sequencing, we built constructs where the endogenous enhancer-promoter-5’UTR for the genes were fused to the coding region of fluorescent Venus protein along with the *PGK1* 3’UTR. The level of expression of Venus was measured using a fluorescent plate reader. The data show a continuum of expression values with the *TDH3* regulatory sequences generating the most fluorescence while the inducible genes produced the least fluorescence in glucose (Figure 8).

Based on these data we then selected regulatory regions from nine of the genes spanning different expression levels. We systematically built a matrix of 81 different constructs where the enhancers of these 9 genes were combined with the promoters and 5’UTRs of these 9 genes. The *PGK1* 3’UTR was used in all of these constructs. Fluorescence was measured using a fluorescent plate reader.

This data matrix can be transformed by normalization to extract information on combinatorial gene regulation. We summed all 81 expression values as the total expression space of the experiment and described the expression of each individual construct as a fraction of this total expression (expressed as a percent of the total). This analysis presents the regulatory strength of each combinatorial cassette as a fraction of the total expression space. This approach allows us to analyze data from different fluorometers with different sensitivities as well as data collected on different days under slightly different growth conditions. Each experiment was repeated between three and seven times (Figure 9).

**Figure 9:**
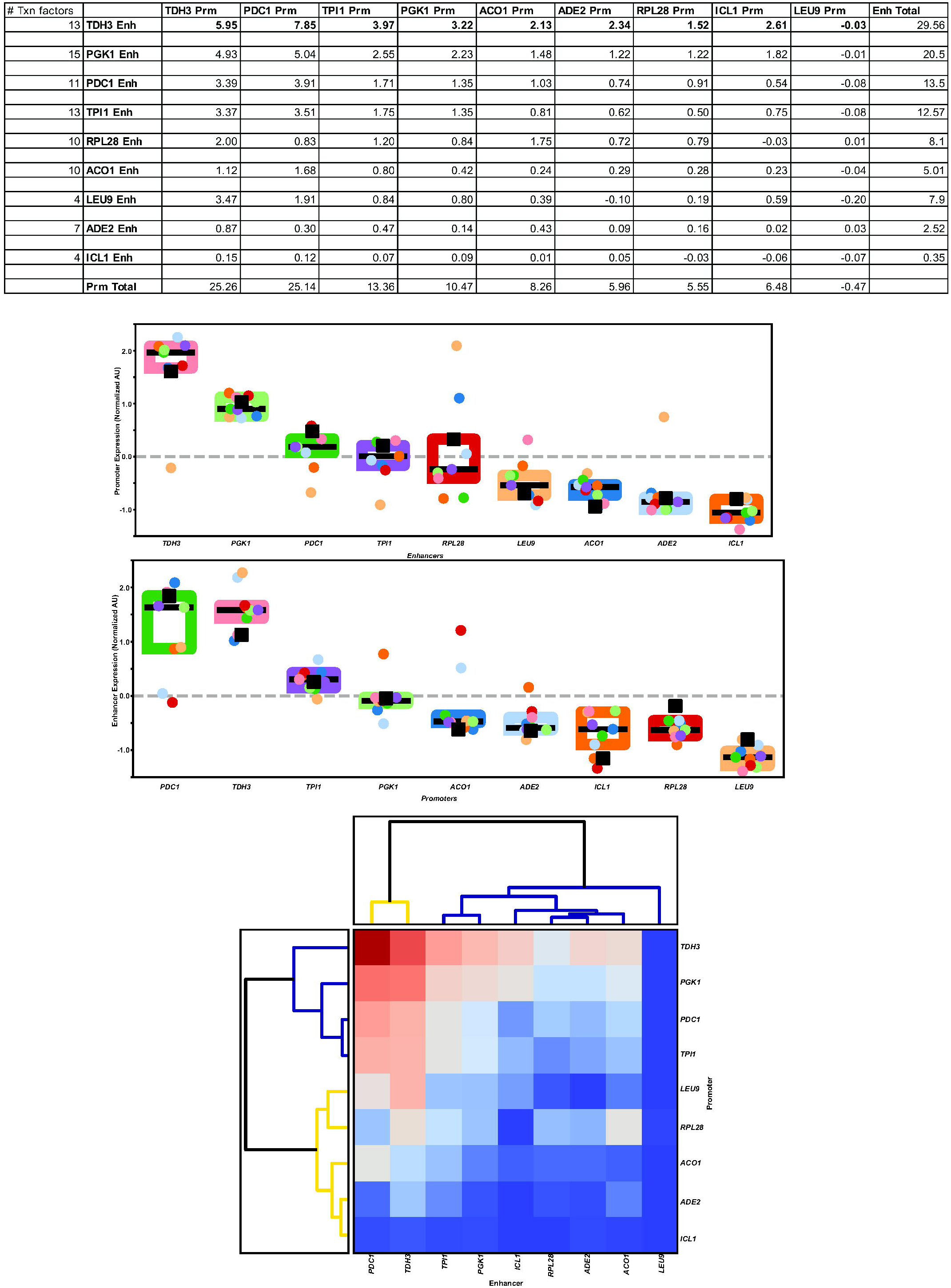
Expression analysis of a 9×9 matrix of Enhancers and promoters+5’UTRs. A 9×9 matrix of different combinations of enhancers and promoters with a Venus reporter gene along with the PGK1 3’UTR was generated. Expression of these constructs was measured using a fluorometer. The expression values of all 81 constructs were summed and the expression of each individual pairwise combination was listed as a percentage of the total expression. Three biological replicates were measured for each construct and the mean values are presented. Box plots of the expression of the 9 enhancers was plotted. Each dot represents the expression level of a specific promoter in combination with that enhancer. Box plots of the expression of the 9 promoters was plotted. Each dot represents the expression level of a specific enhancer in combination with that specific promoter. The Venus expression values of pairwise combinations (of enhancers and promoters) were plotted as a heat map along with clustering analysis to identify relationships between elements.

In glucose containing media, the *PGK1* enhancer with its cognate promoter generates 2.23% of the fluorescence. This value is increased when the *PGK1* enhancer is combined with either the *TDH3* core promoter or the *PDC1* core promoter. Similarly, the *TDH3* enhancer with its cognate promoter generates approximately 5.95% of the total fluorescence landscape. This value increase to 7.85% when the *TDH3* enhancer is combined with the *PDC1* core promoter. Other promoters are unable to increase expression from these strong enhancers and actually reduce expression. For example, the *TDH3* enhancer in combination with the *PGK1* promoter only accounts for 3.22% of the total fluorescence. In comparison to the very strong enhancers, analysis of moderately strong enhancers shows vast increases in expression potential when these enhancers are combined with promoters from other active genes. For example, the *ACO1, RPL28* and *TPI1* enhancers-promoter combinations generate high levels of transcripts but the levels can be increased 2-3-fold by swapping their native promoters with the promoters of the strong genes. These data suggest that while cognate promoters are optimized to work with their native enhancers, optimization does not imply maximizing enhancer mediated expression, there exists an ability to significantly increase expression when these enhancers are combined with other promoters.

Using these data, we determined the enhancer and promoter strengths for all nine genes and plotted these as box plots and as heat maps (Figure 9). Both enhancer and promoter elements positively and negatively influence expression. For example, the native *TPI1* cassette is ranked 3^rd^ in overall expression. However, when the enhancer and promoter are separated, the *TPI1* enhancer is ranked 5^th^ while its promoter is ranked 3^rd^ suggesting that the *TPI1* promoter/5’UTR fragment positively regulates the *TPI1* enhancer. Similarly, the native *LEU9* cassette is ranked 9^th^ in overall expression. However, analyzing its enhancer separated from its native promoter and combined with the promoter of other genes, the *LEU9* enhancer is ranked 6^th^. The *LEU9* promoter divorced from its native enhancer and combined with other gene enhancers is ranked 9^th^, suggesting that the promoter fragment of this gene negatively regulates all enhancers.

### Measurements of expression as a function of different environmental conditions

We took advantage of the 81-cassette matrix to study gene activation by growing the transformants in different growth conditions (Figure 10). We chose galactose containing and glycerol containing media since galactose is a fermentable sugar while glycerol is a non-fermentable sugar. Comparison of the genes grown in glucose, galactose and glycerol containing media (as well as media lacking adenine) shows the role of promoters in integrating signals emanating from the enhancer elements in response to various environmental cues such as a change in carbon source as well as nucleotides. For example, when comparing changes in gene expression in glucose compared to galactose containing media, we find that the *ICL1* enhancer becomes derepressed. Similarly, in glycerol containing media both the *ICL1* and *ACO1* enhancers become active while genes involved in fermentation show reduced activity. The same analysis done in media containing or lacking adenine shows a similar effect for the *ADE2* enhancer.

**Figure 10:**
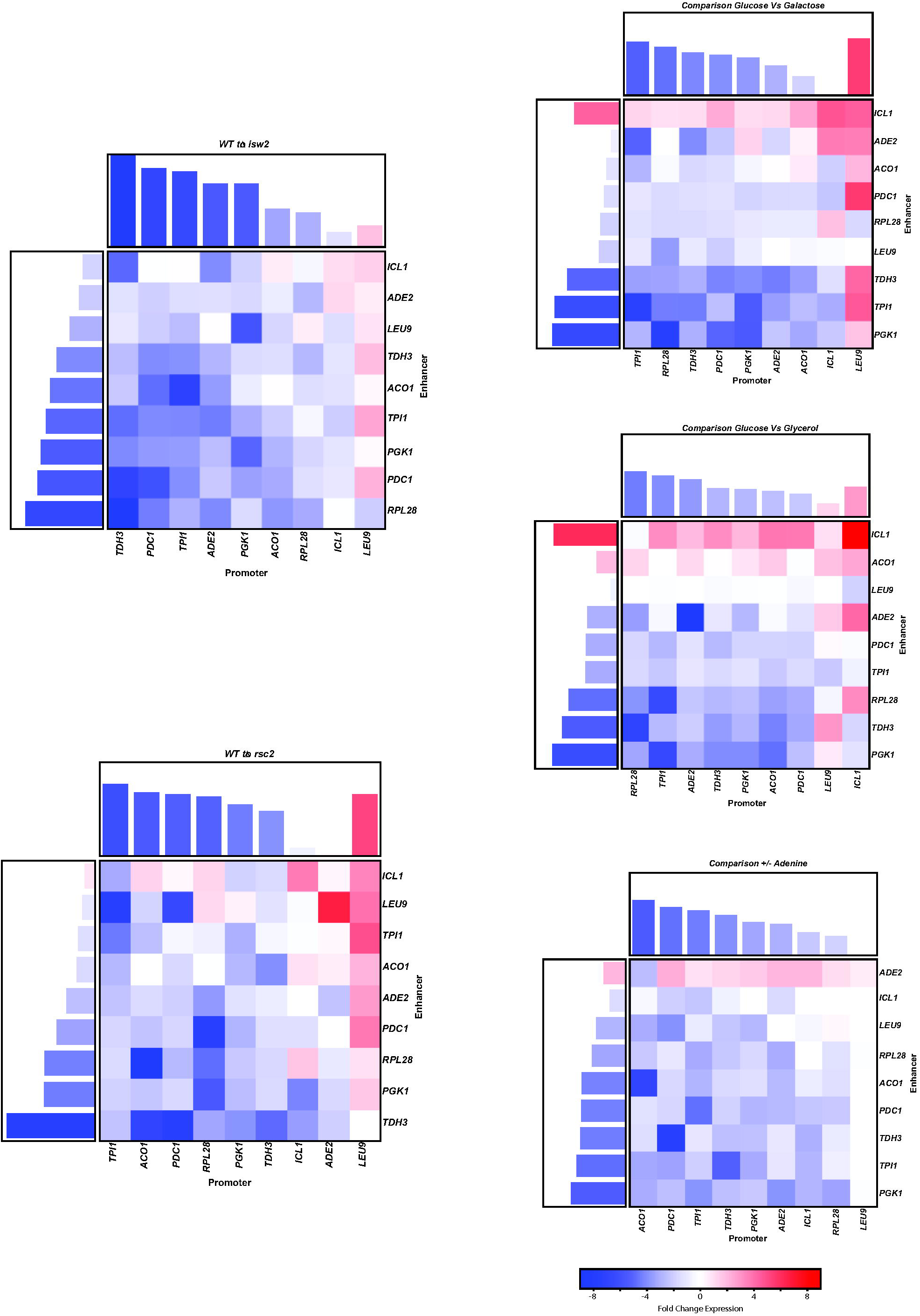
Expression analysis of the 9×9 matrix of enhancers and promoters under varying growth conditions. Cells containing the 81 combinations of enhancers and promoters were grown in different growth conditions and expression of the Venus cassette was measured in a fluorometer. Three biological replicates were measured for each construct. The difference in expression of cells grown in glucose to cells grown in other conditions are plotted as heat maps. Bar graphs above and on the side of the heat map are a summation of the nine individual values.

We also performed this comparative analysis in mutants for chromatin remodeling factors Rsc2 and Isw2. Both of these proteins play a role in organizing the nucleosome free region at the core promoter as well as the −1 and +1 nucleosomes flanking the NFR. In a Rsc2 mutant the fold expression of most genes is reduced though the opposite effect is observed at the *LEU9* promoter suggesting that the repressive effects of the *LEU9* promoter may be Rsc2 dependent possibly via an unfavorable placement of nucleosomes over the core promoter. The same change in expression is observed in an Isw2 mutant. These data seem to suggest that at most promoters, chromatin remodelers are required for gene activation while at the *LEU9* promoter the repressed state is maintained by a specific nucleosome configuration that is weakened in the absence of the chromatin remodeling proteins.

## Discussion

Transcription is a result of a competition between nucleosome binding/dissociation and transcription factor binding/dissociation at the enhancer elements (Lam et al., 2008). The proteins bound to distal enhancer elements stimulate transcription via effective communication with proteins bound to the core promoter. This ultimately manifests itself via the formation of an active transcription complex at the core promoter resulting in transcription. Our data show that changes in the core promoter sequence had similar effects on most enhancers regardless of the arrangement of the transcription factor binding sites within the enhancer. This suggests that one function of the core promoter is to act as a modulator of expression without changing the combinatorial rules involved in the regulation of the gene.

In yeast there are two core promoter architectures-TATA containing core promoters and TATA-less core promoters (Lubliner et al., 2013; Lubliner et al., 2015; Raser and O’Shea, 2004; Smale and Kadonaga, 2003; Vo Ngoc et al., 2017; Zhang and Dietrich, 2005). Our data suggest that the presence of a TATA box can affect the levels of transcripts produced by a enhancer. The underlying molecular mechanism is most likely modulation of TBP binding strength. The presence of a TATA box at a core promoter likely increases the probability of the formation of a functional pre-initiation complex at the promoter since TATA boxes are high affinity binding sites for TBP/TFIID. This is similar to observations in metazoans where some enhancer bound activators (such as E2) require two core promoter elements (TATA box and initiator element) to stably mediate transcription while other activators (such as Sp1) require a single element (TATA box) to stably mediate transcription (Smale, 2001). Thus, weak activators stimulate transcription via a molecular mechanism that benefits from enhanced affinity of TFIID binding to the core promoter while strong activators can mediate high-levels of transcription even in the presence of a sub-optimal core promoter.

In larger eukaryotes there is specificity in communication between enhancer-bound transcription factors and different core promoters. The presence of different core promoter increases enhancer mediated combinatorial control of transcription. For example, the *Drosophila* ribosomal protein gene enhancers activate transcription primarily via interactions with core promoters that are bound by a TBP variant present but are unable to effectively activate promoters bound by TBP. Thus, different forms of core promoters function to restrict the stimulatory ability of nearby enhancers (Butler and Kadonaga, 2001; Kutach and Kadonaga, 2000; Wang et al., 2014). Our principal component analysis of enhancers identified the *RPL28* enhancer as an outlier in its ability to communicate with the different core promoters tested. This exception was also detected in the measurement of gene activity using fluorescent cytometry where the *TDH3* enhancer linked to the *RPL28* promoter did not fall along the universal curve suggesting a cassette specific mechanism of regulation for this construct (data not shown). The *RPL28* gene is part of the ribosomal protein family of genes all of which are coordinately regulated. Our data suggest differences in the mechanism by which this enhancer communicates with core promoters and is consistent with mapping data of transcription factors and chromatin architecture. While genes required for growth during fermentation are regulated in part by the transcription activators Reb1p and Gcr1p, the ribosomal protein family of genes are regulated by Rap1p and Abf1p (Fermi et al., 2016; Reja et al., 2015; Rossi et al., 2018). Reb1p and Gcr1p usually bind near the −1 nucleosome and promote RSC mediated nucleosome mobility immediately downstream of their binding sites (Kubik et al., 2015). Unlike Reb1p, Rap1p typically binds at the −2 and −3 nucleosomes and aids in nucleosome eviction >400bp from its binding site (Fermi et al., 2016; Knight et al., 2014; Kubik et al., 2015; Reja et al., 2015). It is likely that this difference in architecture has effects on transcription from the different core promoters and our combinatorial constructs are able to detect these differences. A better understanding of the mechanisms underlying these differences will require further mutagenic and molecular analysis.

Our data also suggest that some UTRs deserve further analysis as well. The *ICL1* gene is activated in non-fermentable carbon sources. It contains a large 5’UTR whose function is unclear. Our data indicate that this element influences enhancer and promoter mediated gene expression. While the enhancer of the *ICL1* gene has been characterized and contains a CSRE element that represses the gene in the presence of fermentable sugars (Turcotte et al., 2010) nothing is currently known about post-transcriptional regulation of this gene via its UTRs. At other genes, the 5’UTR plays important roles in docking mRNAs to ribosomes and initiating translation. The Kozak sequence from −6 to +6 of the start codon affects protein expression (Dvir et al., 2013) as do sequences further upstream of the Kozak sequence (Cuperus et al., 2017; Hamilton et al., 1987; Li et al., 2017). It will be interesting to know if the long 5’UTR of this gene has additional motifs that regulate this gene at the post-transcriptional level.

The turnover of mRNA is a crucial step in post-transcriptional regulation of genes. This function is primarily mediated by the 3’UTR in a regulated fashion via AU-rich elements. Reports show that these elements can affect protein expression levels significantly (Curran et al., 2013; Yamanishi et al., 2013). We observe a measurable contribution of the 3’UTR/terminator in gene regulation and identify specific 3’UTRs as drivers of expression variation. This effect is specific for certain 3’UTRs since we see the greatest variation in expression with the *ACO1* and *HIS3* fragments though the exact mechanism by which 3’UTRs of these genes affect enhancer and promoter mediated expression is not clear. It will be interesting in the future to determine if this may have something to do with the observation that promoters and terminators of genes are in close 3D proximity in a cell during transcription or whether this is a post-transcriptional effect on mRNA stability and turnover.

Another aim of these experiments was to generate synthetic regulatory elements that exhibited varying activity levels similar to approaches previously used to explore enhancer-promoter combinations (Blazeck and Alper, 2013; Blazeck et al., 2012; Rajkumar et al., 2016). Using this approach, we have identified combinations of regulatory elements that generate a larger spectrum of activity than the native element. These synthetic cassettes, where the same enhancer is coupled with different core promoters, allows one to change expression levels by two orders of magnitude without significantly altering the ability of the cassette to respond to external stimuli and could be a useful resource.

## Material and Methods

### Golden Gate Cloning

All regulatory fragments were PCR amplified from S288c genomic DNA using specific primer pairs. Each fragment was amplified with primers containing a BsmBI recognition site that upon digestion would create sticky ends with the sequences TCGG and GACC at the 5’ and 3’ ends respectively. Adjacent to the BsmBI site, each PCR primer also contained a sequence that is recognized by BsaI and that upon digestion would create specific sticky ends for each regulatory element. Thus, the enhancer fragments had BsaI sticky ends with the sequences AACG and TGGC while the core promoter fragments had BsaI sticky ends with the sequence TGGC and TTCT. The 5’UTR fragments had BsaI sticky ends with the sequence TTCT and TATG, the mRuby2 reporter gene fragment had BsaI sticky ends TATG and ATCC while the 3’UTR fragments had BsaI sticky ends with the sequences ATCC and TCTG. The TGGC sequence in the promoter fragment was followed by the sequence TATGCC. During assembly the 6bp insertion along with the 4bp TGGC Golden Gate scar in effect inserts a 10bp fragment between the upstream enhancer and the TATA box. A 10bp insertion was chosen at this site since 10bp insertions between the enhancer and the TATA box has previously been shown to be optimal for transcription while 5bp insertions are deleterious for optimal transcription (Lubliner et al., 2015).

The amplified DNA fragments were cleaned using a Bioline PCR purification kit. Purified PCR products were quantified using a nanodrop spectrophotometer and cloned into pYTK001 (Lee et al., 2015) with the enzyme BsmBI. 60fmoles of insert were combined with 60 fmoles of plasmid DNA along with 1x T4 DNA ligase buffer, BSA, 1ul BsmBI (NEB) and 1ul high concentration T4 DNA ligase (NEB). The reaction was incubated for 50 cycles at 37C for 3min, and at 16C for 4min. The reaction was terminated by incubating at 50C 5min followed by 80C 5min and stored at 4C until ready to use.

Between 1ul and 2ul of each ligation reaction was transformed into 25ul of DH10B competent cells and plated on 2xTY plates containing chloramphenicol. Between two and five colonies were picked, grown overnight in selective media and plasmid was isolated using a Qiagen mini-plasmid purification kit. Plasmids were checked for inserts using insert specific primers.

The 103 parts plasmids were then used to create a combinatorial library such that the different fragments would combine in the correct order but in a random manner. To join the different regulatory fragments, we mixed 30 fmoles of each parts plasmid containing the different regulatory fragments with 30 fmoles of vector (a *ARS/CEN/URA3* derivative of pYTK096 (Lee et al., 2015)) in the presence of the enzyme BsaI in a Golden Gate reaction as described above. Multiple reactions were performed in parallel and pooled prior to E. coli transformation. DH10B competent cells were electroporated with the library and transformed cells were plated on multiple large 2xTY plates containing Kanamycin. Cells from the plates were scraped off the plates, and plasmid DNA was isolated from these cells and purified on a Cesium Chloride gradient.

### Yeast Electroporation of Library

Yeast strain (ROY5634: MATa ADE2 lys2D leu2-3,112 his3-11 ura3-1 trp1-1 can1-100 rpl18b::BFP2-2) was transformed with the library DNA (Meilhoc et al., 1990). Cells were grown overnight in 5ml YPD medium. 200ml fresh YPD was inoculated with the overnight culture at a concentration of 0.25 OD/ml and grown for 5h. Cells were spun and resuspended in 50ml YPD with Tris:DTT and incubated at 30C for 30min with shaking. Cells were washed with 25ml Buffer-E (10mM Tris HCl pH 7.5, 270mM Sucrose, 1mM MgCl_2_) and resuspended in 2ml Buffer-E DNA was added to the cells and 100ul of cells were placed in a 0.2mm cuvette and electroporated at 540V 25uF, infinite resistance with an exponential pulse. Cells were resuspended with YPD, incubated at 30C for 1h without shaking and then plated on YMD plates lacking uracil. After 3 days, approximately 10,000 colonies were present on each plate. Cells were scrapped off the plate into YMD lacking uracil media, grown for 5h at 30C and then frozen at −70C in the presence of 20% glycerol.

### Cell sorting

5ml of yeast cells containing the library were used to inoculate 50ml YMD medium containing adenine, leucine, lysine, histidine and tryptophan (lacking uracil) and cells were grown at room temperature overnight (approximately 3 doublings). This culture was then used to inoculate 250ml YMD-uracil and grown at 30C for 3h. Cells were pelleted and resuspended into 250ul 1xPBS and 1%BSA at a concentration of 1 OD/ml, filtered through a Nitex mesh and sorted.

### Insert library preps from sorted cells

Total DNA was prepared from the unsorted library, no expression sorted fraction (2.36×10^7^ cells), low expressing sorted fraction (19.5×10^6^ cells), medium expressing sorted fraction (17.1×10^6^ cells) and high expressing sorted fraction (6×10^6^ cells) using the YeaStar Genomic DNA kit from ZymoResearch. 200ng of Nanodrop quantitated DNA from each fraction were treated with ExoVII and Exo VIII (Truncated) (NEB) to reduce the amount of linear genomic DNA. Following denaturation of the enzymes at 80C for 10’, inserts in the plasmid library fractions were amplified with primers using the KAPA HiFi PCR kit in 25ul reactions (10x-50x). Reactions were pooled and precipitated with glycogen and ethanol and the precipitated DNA was re-suspended in nuclease free water. AmPure XP beads (0.8 vol) were used to purify DNA >700bp from the sorted insert libraries. This DNA was quantitated using the Qubit Broad Range DNA kit and subsequently analyzed by conventional agarose and BioAnalyzer gel electrophoresis and eventually used for sequencing.

### Oxford Nanopore Sequencing

700ng of DNA from each of the no, low, medium and high expressing sorted insert libraries were used to prepare samples for sequencing on the Oxford Nanopore MinION. Nanopore barcodes (NB01 > NB04) were individually ligated to each end-repaired and dA tailed fraction, which was quantitated and pooled in the following ratios prior to adapter ligation:

|  |  |  |
| --- | --- | --- |
| Zero: | 63.8% | 448ng |
| Low: | 26.3% | 182ng |
| Medium: | 7.7% | 56ng |
| High: | 2.14% | 14ng |
| Total: | 100% | 700ng |

Adapters were ligated to the pooled barcoded libraries according to the Oxford Nanopore protocol. The DNA was then quantitated and loaded on the nanopore flow cell for sequencing.

In a second sequencing round, libraries were barcoded as before but pooled in the following ratios:

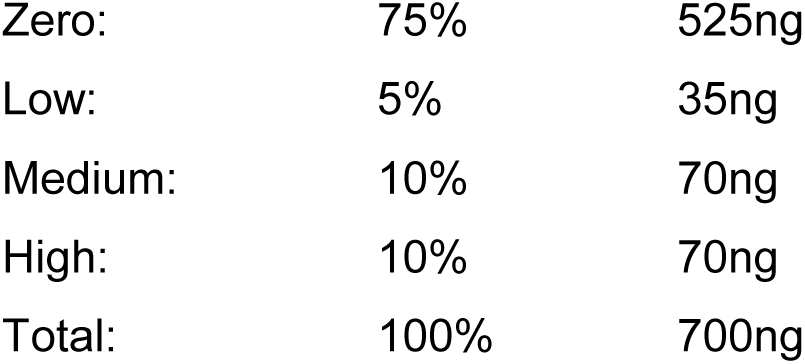

In a third round, 700ng of DNA from the unsorted library were sequenced on the Nanopore sequencing platform. Barcodes were not used in this protocol since the DNA library was from one source.

### Microtiter Plate Transformations of Yeast Cells

Yeast strains were grown for 6h at 30C in 5ml YPD with shaking. 400ml YPD was inoculated with these cells so that the final concentration of cells after 14h of growth is 2 OD/ml. 300ml of cells were pelleted, washed in 150ml 0.1M lithium acetate and resuspended in 3ml 0.1M lithium acetate. 333ul sonicated salmon sperm DNA (10mg/ml) was added to the cells. In each well of a microtiter plate 3-5ul plasmid DNA was added. 25ul of yeast cells were mixed with the plasmid DNAs in the microtiter plates and the cells were incubated at 30C for 15min. 150ul PEG/LiAc was added to each well and mixed by pipetting. The cells were further incubated for 30min at 30C. 17ul DMSO was added to the cells and the plates were heat shocked at 42C for 15min in a heat block. Cells were spun and supernatant was aspirated off. Cells were resuspended in 10ul water and plated onto plates (YMD lacking uracil). Colonies were allowed to grow for 3 days at 30C (Winzeler et al., 1999).

### Fluorescence Measurements using a Plate Reader

Transformed yeast cells were transferred from a plate using a frogging tool (Sigma) into three different microtiter plates each containing 100ul YMD-U media. Cells were grown overnight at 30C without shaking. 30ul of each culture was used to inoculate deep well (2ml) microtiter plates containing 570ul YMD-U. Plates were inoculated overnight at 30C with shaking at 600rpm. 30ul of these overnight cultures were used to inoculate fresh 2ml microtiter plates containing 570ul YMD-U and grown overnight at 30C with shaking. 100ul of fresh YMD-U media in 2ml microtiter plates were inoculated with 50ul overnight cultures and grown at 30C for 3h with shaking. 100ul of each culture was removed and fluorescence was measured using a microtiter fluorescent plate reader.

### Determination of Expression in sorted yeast cells

To determine the mean and variance of expression for a regulatory fragment we fit an estimate of that fragment’s prevalence in each fraction to a log-normal model of protein expression, as described (Townshend et al., 2015). The estimate, *x_i,b_*, of the ratio of cells containing fragment, *i*, sorted into each fraction, *b*, was determined by normalizing the number of reads, *r_i,b_* by multiplication with 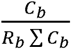, where *C_b_* and *R_b_* are the total number of cells sorted and reads mapped from bin *b*, respectively (calculating the fractional representation of fragment, *i*, in bin *b* and subsequently scaling that fraction by the fraction of cells observed in bin *b* by FACS). We then assume that *x_i,b_* are random variables sampled from binned log-normal distributions where the bins are determined by the FACS fraction boundaries.

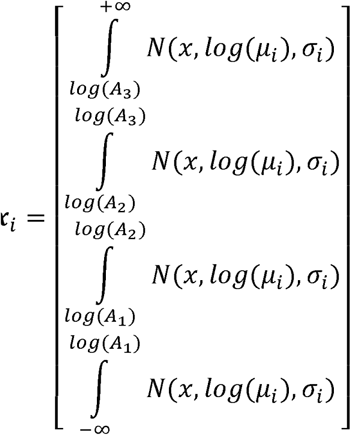

Where, *x_i_*, is the vector of ratios for all bins described above, *µ_i_*, is the mean expression, *σ_i_*, is the standard deviation of expression, *A_b_*, is the expression value for the upper boundary of bin b by FACS.

Supplementary Figure 1: Plots of nucleosomes and transcription factor distribution at the 26 genes

## Acknowledgements

This work was supported in part by a grant from the NIH to RTK (GM078068). M.J. was supported on an NIH grant to M. Akeson (HG007827).

