## Supplementary Figure1 for "Combinatorial analysis of *Saccharomyces cerevisiae* regulatory elements"

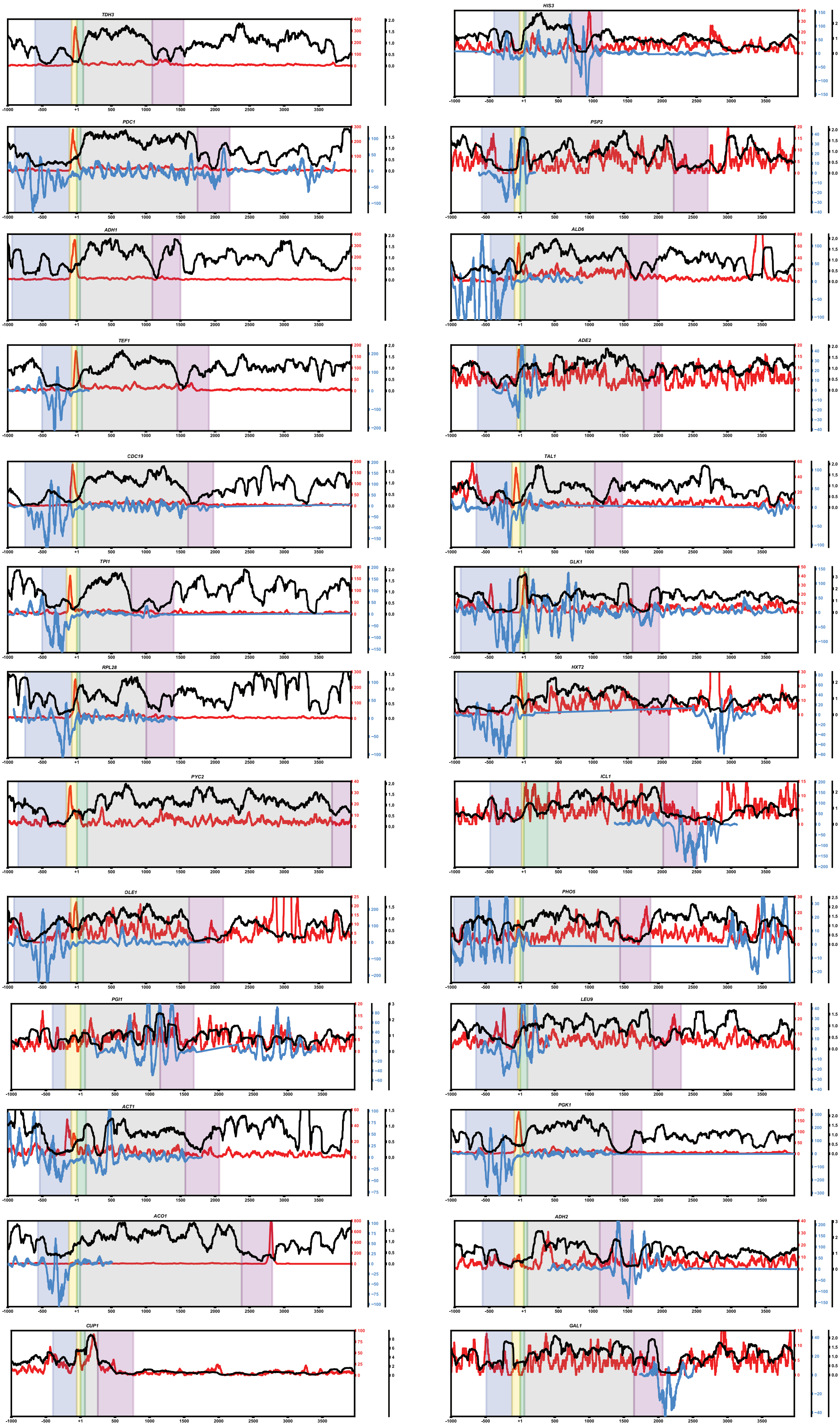


Blue is ATAC seq accessibility data
Red is TBP ChIP-Seq data
Black is Micrococcal-seq nucleosome mapping data
